# Tuning a light-regulated allosteric switch for enhanced temporal control of protein activity

**DOI:** 10.64898/2026.04.01.715907

**Authors:** Jacob Matsche, Jordan Fauser, Trisha Bansal, Nicholas Leschinsky, Courtney Coleman, Andrei V. Karginov

## Abstract

Optogenetics enables researchers to control protein localization, interactions, and activity using photosensitive domains. The key desired properties for optogenetic tools include broad applicability, tight light-regulated control with high dynamic range, and tunability. Previously, we described an engineered light-sensitive switch, LightR, composed of two VVD domains connected by a flexible linker, enabling light-dependent allosteric control of protein activity through site-specific insertion. Here, we introduce enhanced LightR variants with improved dynamic range and faster activation kinetics. Through targeted modifications to the VVD domains and linker region, we optimized a LightR-regulated Src kinase (LightR-Src) activity and generated two LightR-Src variants: one supporting sustained Src activation comparable to constitutively active Src, and another enabling rapid, reversible control, ideal for modeling transient signaling events suitable to mimic Src signaling in living cells. These modifications expand the versatility of LightR-based tools, facilitating their use in diverse optogenetic applications requiring high dynamic range of regulation and fast control of targeted proteins.

## Introduction

Optogenetic tools provide unique capabilities for many areas of biological research that range from dissection of a protein’s function to construction of synthetic signaling circuits regulated by light. Commonly used methods employ homo- and heterodimerizing photosensitive domains for light-mediated control of protein localization, interactions, and oligomerization. These tools demonstrate broad applicability due to simple modular design, tunability, and simple application principles ^1–7^. However, the application of light-sensitive domains for regulation of protein activity has been more challenging as it often requires development of custom approaches suitable only for one or a limited number of proteins and/or exhibits additional constraints. Reconstitution of split proteins by photo-regulated dimerization has been used for regulation of several enzymes but this approach is applicable to a limited number of proteins and requires equimolar expression of two constructs ^8–10^. Steric or allosteric regulation of protein activity using engineered photosensitive domains overcomes some limitations of split protein system as it requires expression of only one engineered protein ^11–13^. However, many of these approaches are limited to a specific protein or specific class of proteins ^10–12,14–16^. Furthermore, these approaches often have limited tunability and thus do not allow researchers to achieve the desired dynamic range of regulation necessary for a specific task. The ideal optogenetic approach for allosteric regulation of protein function should provide flexibility in making adjustments that will ensure broad applicability of the tool.

We have recently developed an optogenetic approach that achieves allosteric regulation of protein function and can be applied to different enzymes ^17^. We engineered a light-regulated switch domain, LightR, composed of two light-sensitive VVD domains tandemly connected by a flexible linker. VVD domains dimerize upon blue light illumination and dissociate in the dark, creating a clamp-like switch that is open in the dark and closed in the light. When inserted into the functional domain of a protein of interest, the open LightR domain will cause allosteric disruption of protein function in the dark. Upon blue light illumination, the LightR domain closes recovering the structure of the targeted protein and rescuing its activity (**Figure 1A**). We also demonstrated tunability of the LightR switch by creating two variants. One construct, named LightR, shows robust activation upon illumination but slow inactivation. LightR domain is suitable for experiments that require prolonged activation that can be maintained by infrequent illumination thus avoiding potential unwanted phototoxicity caused by prolonged illumination. Another construct, FastLightR, demonstrates fast inactivation kinetics enabling cyclical activation/inactivation of a protein and local subcellular regulation in living cells. We successfully applied this tool to achieve regulation of several protein kinases and a DNA recombinase Cre. Despite multiple advantages of the tool, we observed that some proteins regulated by LightR and FastLightR exhibited reduced activity when compared to constitutively active counterparts ^17^.

**Figure 1:**
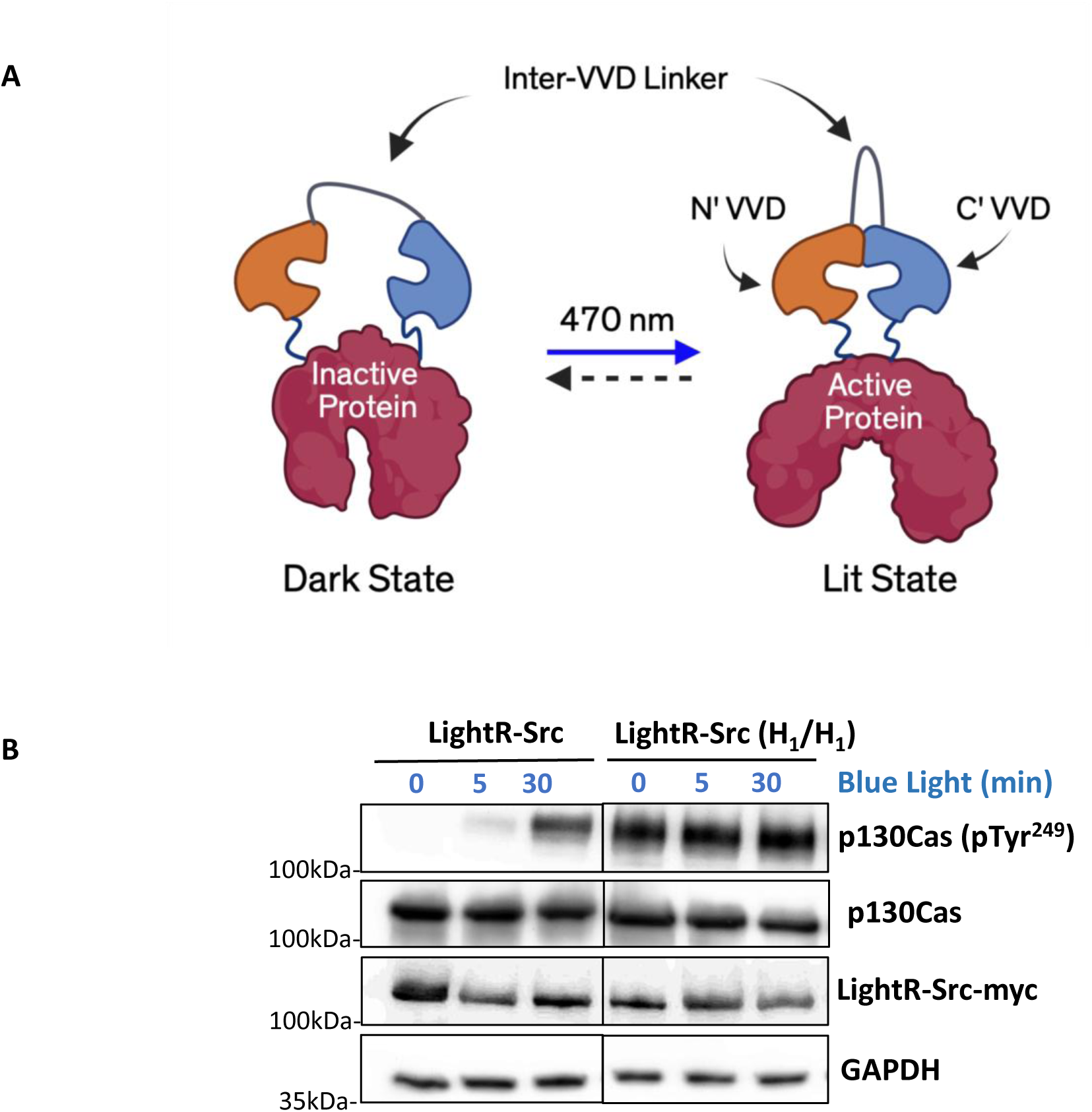
M135I/M165I mutations in LightR switch disrupt regulation of activity by light. (A) Schematic of light-mediated allosteric regulation of proteins by LightR domain comprised of two tandemly connected VVD proteins. (B) 293T cells transiently expressing the indicated Src constructs bearing mCherry and myc tags at the C-terminus were continuously exposed to light for the specified times. Cell lysates were collected and probed for phosphorylation of Src substrate, p130Cas. A representative result of three independent experiments is shown.

Thus, we set out to develop protein engineering strategies that will expand the dynamic range of LightR regulation enabling broader application of this tool.

Here we describe a novel approach for tuning LightR-mediated regulation of protein function that overcomes previous limitations and significantly expands the dynamic range of regulation. Using regulation of tyrosine kinase Src as a model system, we demonstrate that mutations withing VVD domains of LightR allow us to achieve higher activity of LightR-Src in the lit state. Development of a novel linker connecting the two VVD domains enabled efficient elimination of leaky activity in the dark for modified LightR domains. Using our engineering approach, we generated slow- and fast-inactivating versions of light-regulated Src. These versions display significantly higher activity when illuminated with blue light while retaining tight control of activity in the dark. The new method simplifies application of LightR constructs in living cells and provides more flexibility for optimization when employed for regulation of new proteins.

## Results

### Stabilization of the lit state of LightR domain causes leaky activity in the dark

Our previous studies demonstrated that LightR-Src demonstrates robust activation, showing significant increase in phosphorylation of substrates as early as 10 seconds after start of illumination. However, we noticed that fully activated LightR-Src still has significantly lower activity than constitutively active Src mutant (equivalent of Y527F mutation in avian Src) (**Supplementary Figure 1**) ^17^. Thus, we sought for a solution that will allow us to improve activation of LightR-Src and significantly expand the dynamic range of optogenetic regulation by LightR switch. Development of such an approach will have multiple benefits. It will allow researchers to express lower amounts of the constructs while still achieving the desired level of regulation. It will also broaden application of this strategy for regulation of other proteins where original LightR switch may not provide sufficient activation.

To achieve stronger activation of LightR-Src we decided to implement mutations that will stabilize the dimerized state of VVDs upon illumination. Previous studies demonstrated that two mutations (M135I/M165I) can achieve this through stabilization of the cysteine-flavin adduct in the photosensitive core of the protein ^4^. Indeed, LightR-Src construct bearing M135I/M165I in both VVD domains (LightR(H_1_/H_1_) demonstrated higher activity than the original LightR-Src but remained active in the dark (**Figure 1B**). This result suggests that stabilization of the lit state of VVD favors closed conformation of LightR switch and thus makes it active even in the dark.

### Modification of the flexible inter-VVD linker in the LightR switch enhances dynamic range of LightR-Src regulation

Unlike free floating VVD monomers, the proximity of the VVD domains within LightR increases the likelihood of VVD dimerization in the dark state. Thus, by stabilizing the dimer interaction with the M135I/M165I mutations, we potentially generated a LightR-Src construct that exists in the on-state even in the dark. Original LightR domain utilized a long flexible linker composed of Gly- Ser-Gly (GSG) repeats to connect the two VVD monomers (inter-VVD linker) (**Figure 1A**). This flexibility was intentionally designed to accommodate the conformational changes exhibited by the LightR switch as it transitioned between the dark and lit states. However, in the case of stabilizing M135I/M165I mutations, high flexibility of the linker may allow for interaction between VVD domains in the dark creating a leaky switch. We hypothesized that by modifying the flexibility of the inter-VVD linker we can eliminate leaky activity of LightR-Src (H_1_/H_1_) in the dark while retaining elevated activity upon illumination. Application of a more rigid inter-VVD linker may force an open conformation of the LightR switch in the dark thus preventing unwanted leaky activity.

We designed several different inter-VVD linkers with varying degrees of rigidity: Spider-Silk linker (SSL) derived from flagelliform protein, Ferredoxin-Like linker (sFL, a fragment of ferredoxin-like domain), and Worm-Like Chain Linker (WLC) (**Figure 2A**). They can transition between structured and flexible unfolded conformations^18–20^. The structured conformation will provide increased rigidity that could reduce leaky dimerization of the VVD domains in the LightR switch in the dark. On the other hand, more flexible unfolded conformation should allow for light-induced closing of the LightR. SSL linker is predicted to have spring-like property capable of compressing and relaxing ^21,22^. sFL linker may exhibit a helical organization that establishes electrostatic interactions stabilizing a specific rigid conformation, as suggested by molecular dynamics simulations (**Supplementary Figure 2**). The Worm-like-chain linker is a spontaneously collapsing helix. This linker primarily exists in an extended conformation, however due to the alternating charged residues throughout the helix, it can bend into a conformation which brings the termini closer to one another ^20^. We hypothesized that testing the linkers with variable degree of rigidity will allow us to identify constructs that would eliminate leaky dark state activity of LightR-Src H_1_/H_1_, but support efficient activation upon stimulation with blue light.

**Figure 2:**
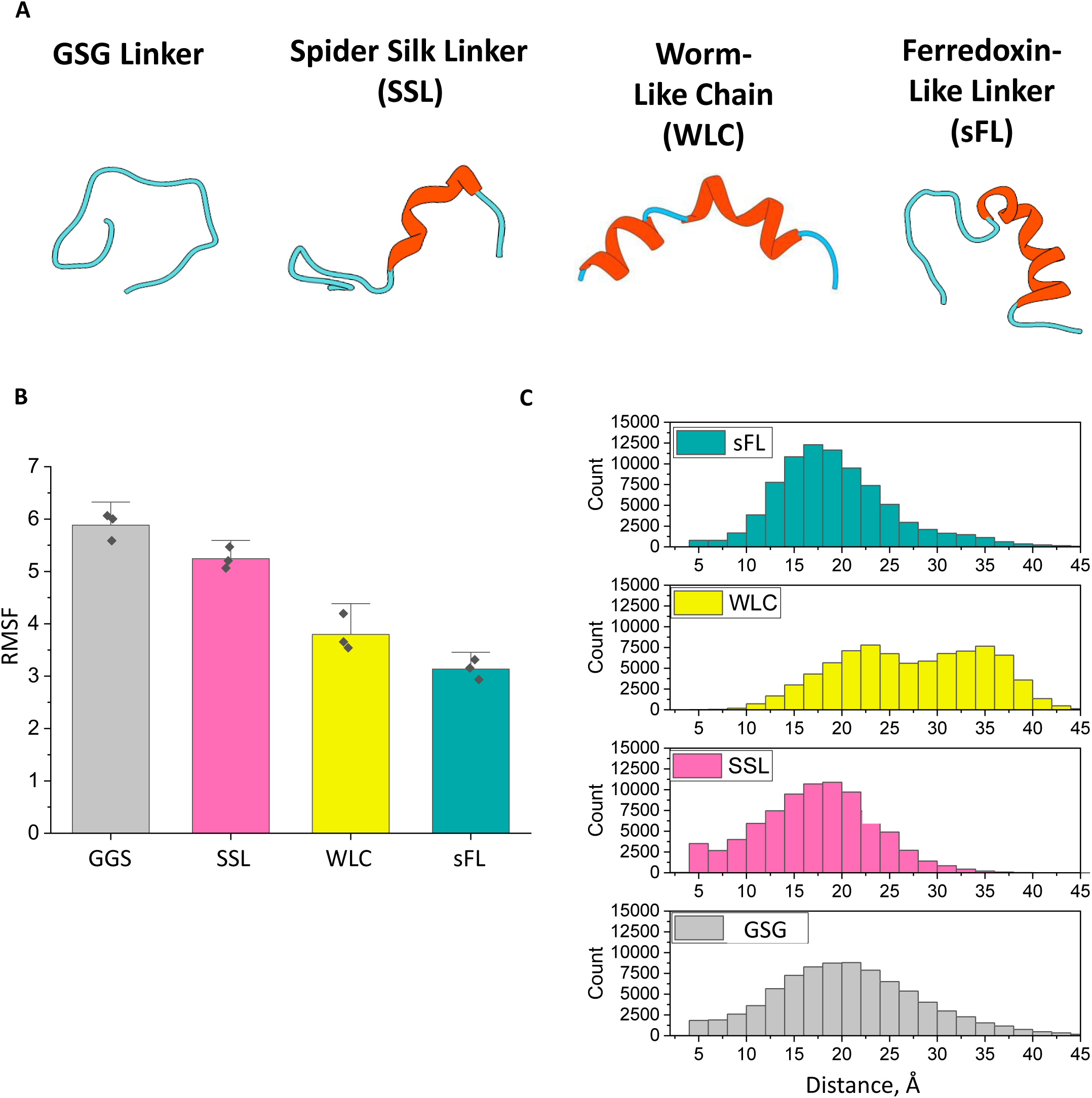
Molecular modeling of different inter-VVD linkers. (A) Representative models of selected linkers with unstructured regions in blue and structured regions in orange. All models were generated, minimized, and equilibrated using Chimera. (B-C) Molecular dynamics simulations were carried out at 310 K in an implicit solvent field using NAMD. (B) RMSF of the linkers determined for three 45 ns simulations. (C) Distribution of distances between the Cα of the N- and C-termini of each linker for 82,500 conformations.

Molecular dynamics simulations of these linkers confirmed that they exhibit variable degrees rigidity. Evaluation of the Root Mean Square Fluctuation (RMSF) of each linker revealed a gradient of flexibility and dynamics within the linkers (**Figure 2B**). The original (GSG) and spider-silk (SSL) linkers exhibit a higher degree of flexibility compared to the worm-like chain (WLC) or ferredoxin-like (sFL) linkers. Further analysis of the linker dynamics provided information on the likelihood of the N- and C-termini of the linker coming in proximity to one another, which could potentially influence leaky activity in the dark (**Figure 2C**). The original GSG linker exhibited a broad range of distances between the C- and N-termini reflective of its high flexibility. The SSL and sFL linkers demonstrated a tighter range of distances between the ends due to their increased rigidity. Notably, the SSL linker showed a shift to shorter distances between the ends suggesting that it mayfavor a closed conformation of the LightR switch. The sFL linker exhibited a lower probability of allowing shorter distances (5-10 Å) which suggests the linker may act as a more rigid spacer between the VVD monomers (**Figure 2C**). Further evaluation of the hydrogen bonding network within the sFL linker revealed two Arg residues (Arg^8^ and Arg^25^) form several hydrogen bonds with the backbone of the opposing terminus (**Supplementary Figure 2**). These likely contribute to the reduced conformational sampling of this linker, and due to the size of the Arg residues provide a sufficiently spaced rigid linker between the two VVDs. Use of this linker as a rigid spacer between the two VVDs may enable a reduction in leaky dark state activity, however this may also come at the cost of robust activity in weakly activated constructs. Interestingly, the Worm-like chain linker exhibited a biphasic distribution, suggesting it can sample both extended and contracted conformations and may facilitate more favorable transitions between the lit and dark state (**Figure 2C**). However, this linker shows limited sampling of conformations with the termini under 12 Å apart. This may result in reduced activity in the lit state with WLC linker due to increased disruption of the lit state dimer.

Based upon the molecular dynamics data suggesting the potential of the new inter-VVD linkers act as a rigid spacer and disrupt dark state dimerization, we implemented each of the new inter-VVD linkers into LightR-Src (H_1_/H_1_). Increasing rigidity of the linker resulted in reduction of activity in the dark state, with both *WLC*-LightR-Src (H_1_/H_1_) and s*FL*-LightR-Src (H_1_/H_1_) exhibiting improved dynamic range, while *SSL*-LightR-Src (H_1_/H_1_) did not significantly improve the leaky activity in the dark (**Figure 3A**, **Supplementary Figure 3A**). Our results show that s*FL-*LightR-Src (H_1_/H_1_) exhibited the optimal dynamic range among these new constructs, with robust activity in the lit state and minimal activity in the dark state. This improved construct was named HiLightR-Src and was selected for further evaluation.

**Figure 3.**
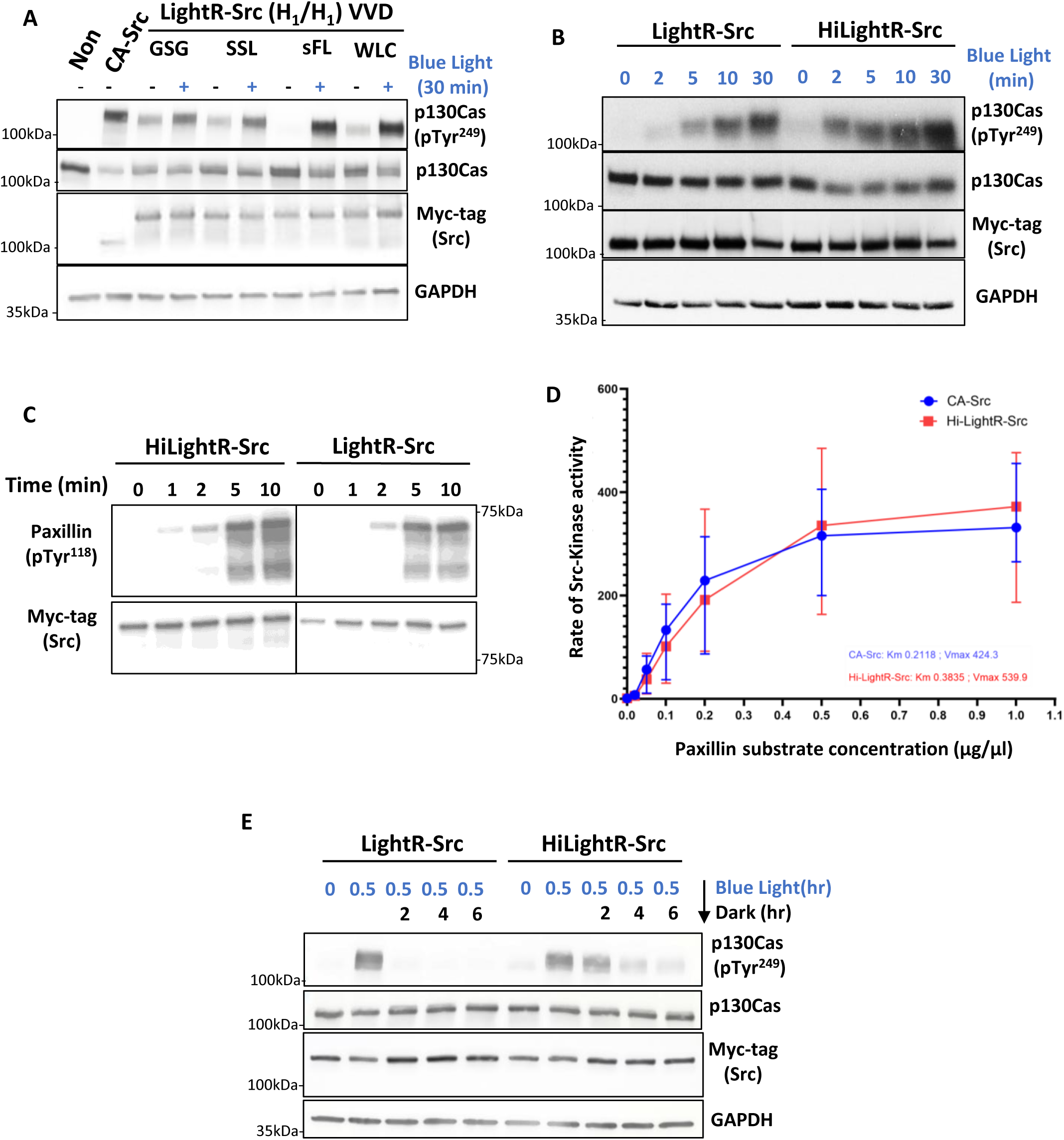
sFL inter-VVD linker in combination with M135I/M165I mutations improve activation of LightR-Src. (A-B), Phosphorylation of endogenous p130Cas in response to activation of indicated Src constructs in living cells. (A) HEK293T cells transiently expressing the indicated LightR-Src (H1/H1) constructs, each bearing indicated inter-VVD linker and tagged with mCherry and myc-tag at the C-terminus, were continuously exposed to blue light for indicated periods of time. Cell lysates were collected and probed for phosphorylation of Src substrate, p130Cas, via Western Blot. (B) HEK293T cells transiently expressing the indicated Src constructs bearing mCherry and myc tags at the C-terminus were continuously exposed to light for the specified times. Cell lysates were collected and probed for phosphorylation of Src substrate p130Cas. (C) Analysis of HiLightR-Src activity using *in vitro* assay. HEK293T cells transiently expressing the indicated Src constructs bearing mCherry and myc tags at the C-terminus were exposed to continuous blue light for 30 minutes. Src constructs were then immunoprecipitated and their ability to phosphorylate purified N-terminal fragment of paxillin was assessed. Src constructs and fragment of paxillin were incubated at 37℃ for indicated amounts of time before analyzing by western blot. (D) Km and Vmax HiLightR-Src and CA-Src determined by *in vitro* kinase assay by varying substrate concentration. Analysis graphs are shown as mean ± 95% confidence interval, N=3. (E) Inactivation kinetics of LightR-Src or HiLightR-Src. HEK293T cells transiently expressing the indicated Src constructs bearing mCherry and myc tags at the C-terminus were continuously exposed to blue light for 30 minutes. Cells, where dark time periods are indicated, were subsequently incubated in the dark for the specified durations following light illumination. Cell lysates were collected and probed for phosphorylation of p130Cas. All experiments were repeated at least three times with similar results.

To evaluate the improved capabilities of HiLightR-Src we compared its activation kinetics and levels of activity to the original LightR-Src construct. HiLightR-Src exhibited significantly faster activation kinetics with significantly higher levels of activity after two minutes of illumination (**Figure 3B**, **Supplementary Figure 3B**). In vitro kinase assay analysis also demonstrated that HiLightR-Src has higher activity than LightR-Src at levels comparable to constitutively active Src (**Figure 3C**, **D**). Inactivation of HiLightR-Src after placement of cells in the dark was significantly slower than LightR-Src requiring greater than 4 hours to fully deactivate (**Figure 3E**, **Supplementary Figure 3C**). This property of HiLightR-Src will be advantageous for limiting phototoxicity in long term activation experiments as it will require brief exposure to light to retain robust activity for extended periods of time. Overall, application of a novel linker with increased rigidity allowed us to generate an improved HiLightR-Src construct that demonstrates greater dynamic range of activation, faster activation kinetics, and prolonged activity following stimulation.

### Stabilizing mutations in VVD significantly improves activity of FastLightR-Src

Slowly inactivating LightR constructs are useful for studies where prolonged activation is desired. However, projects focused on dissection of fast transient signaling and/or requiring local subcellular regulation of a protein need light-regulated switched demonstrating fast inactivation. Our previous work described the generation of FastLightR switch that demonstrates fast inactivation kinetics and enables subcellular control of protein activity ^17^. However, FastLightR-Src exhibited significantly reduced activity limiting applicability of the tool ^17^(**Figure 4A**). We hypothesized that stabilizing VVD dimer in the lit conformation may improve activity level of FastLightR-Src. To achieve that, we introduced lit-state stabilizing M135I/M165I mutation in one of the VVD domains within FastLightR generating FastLightR(H_1_)-Src. However, this modification did not significantly increase activity upon illumination with blue light and led to much slower inactivation kinetics (**Figure 4B**)

**Figure 4:**
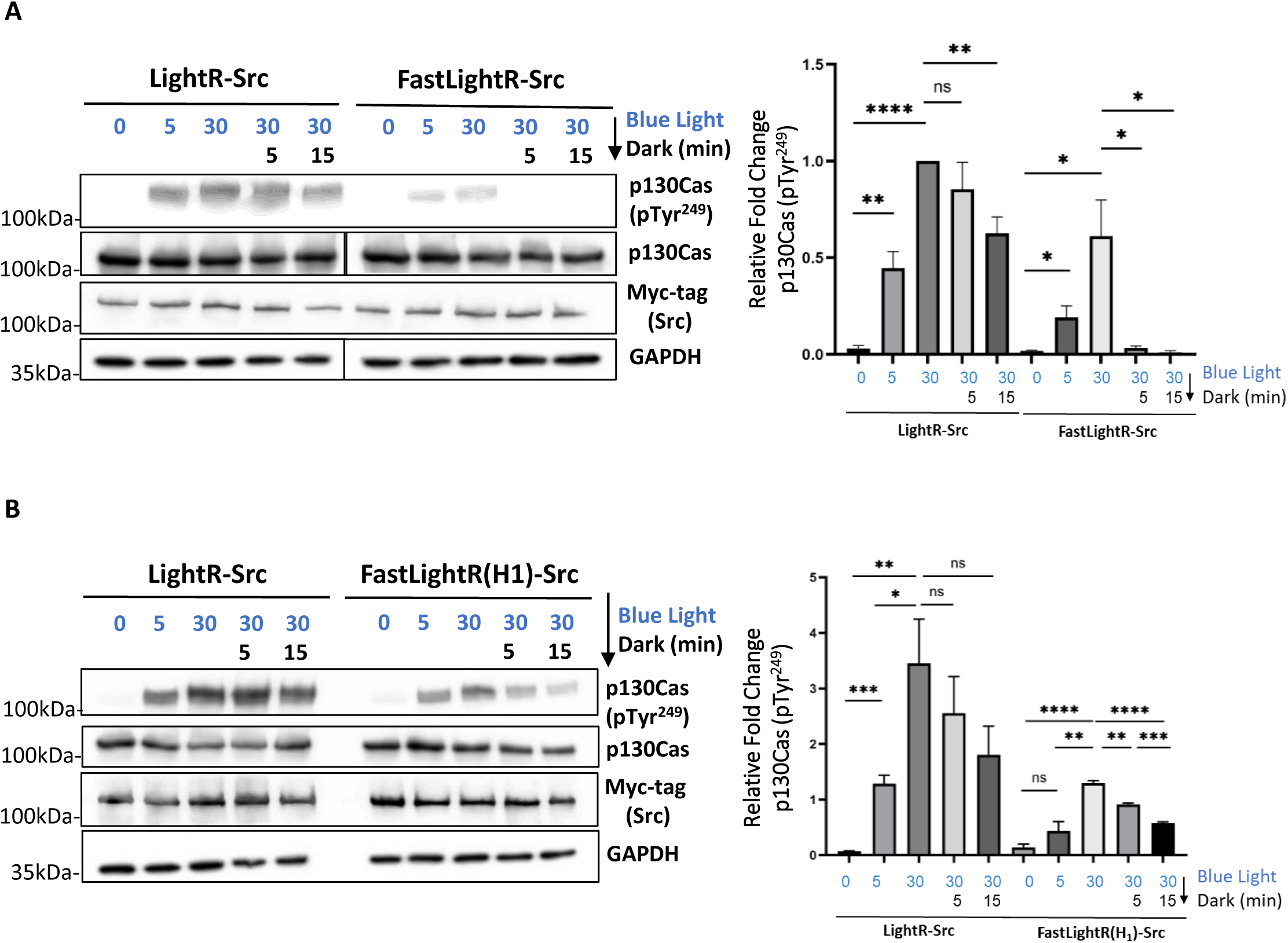
M135I/M165I mutations in VVD do not improve activity of FastLightR-Src. (A, B) HEK293T cells transiently expressing the indicated Src constructs bearing mCherry and myc tags at the C-terminus were continuously exposed to light for the specified times. Cells, where dark time periods are indicated, were then placed in the dark for different periods of time. Cell lysates were collected and probed for phosphorylation of Src substrates. All experiments were repeated at least three times. Quantification of p130Cas phosphorylation is shown as mean ± standard errors of mean (SEM) and statistical differences were examined by unpaired Student’s t-test. NS: not significant (p>0.05) ; * : significant (p<0.05) ; *** : significant (p<0.001) ; **** : significant (p<0.0001).

A recent study described a series of mutations in modified VVD domains (eMagnets) that significantly improve their application for light-regulated dimerization of proteins ^1^. These mutations improve the stability of the protein and its dimerization efficiency in cultured mammalian cells. Importantly, these constructs demonstrate fast kinetics of dimerization and dissociation. Thus, we decided to employ these mutations to improve regulation of Src by FastLightR. We introduced mutations (T69L, Y94E, S99A, N100R, A101H, N133Y or N133F, R136K, and M179I) in both VVD domains of the LightR switch of FastLightR-Src to generate two constructs, with either the N133Y and N133F mutations (N133Y-eFastLightR and N133F-eFastLightR respectively). Previous studies demonstrated that N133Y mutation exhibits stronger dimerization of the VVD domains while the N133F mutation displayed faster dissociation kinetics^23^. Both N133Y and N133F eFastLightR-Src mutants exhibited higher activity levels in the lit state compared to FastLightR-Src counterpart (**Figure 5A**, **Supplementary Figure 4A**). However, both also exhibited leaky activity in the dark state. The N133F mutant demonstrated less leakiness in the dark, which correlated with the suggested weaker dimerization efficiency of that mutant when compared to the N133Y mutant. To reduce the leaky activity of these constructs, we modified N133Y- and N133F-eFastLightR-Src by replacing the GSG linker connecting the two VVD domains with the sFL linker. The addition of the sFL linker dramatically reduced the observed leaky activity in the dark (**Figure 5B**, **Supplementary Figure 4B**). Since N133F-eFastLightR-Src with sFL linker demonstrated strongest activation and no leaky activity, we selected this construct for further analysis and named it eFastLightR-Src. Next, we compared the activation and deactivation kinetics of eFastLightR-Src with our previously published FastLightR-Src construct. Our analysis showed that eFastLightR-Src exhibits significantly faster activation (**Figure 5C**, **Supplementary Figure 4C**). Importantly, eFastLightR-Src demonstrated comparable inactivation kinetics in living cells (**Figure 5D**, **Supplementary Figure 4D**). Although phosphorylation levels of paxillin and p130Cas were still higher at 5 minutes following inactivation, this could be in part attributed to higher phosphorylation level before cells were placed in the dark (**Figure 5D**, **Supplementary Figure 4D**). Repeated cycles of activation/inactivation demonstrated that eFastLightR-Src consistently retains higher activity when illuminated and inactivates down to basal level in the dark, enabling more efficient cyclical regulation of Src activity in living cells (**Figure 5E**, **Supplementary Figure 4E).**

**Figure 5:**
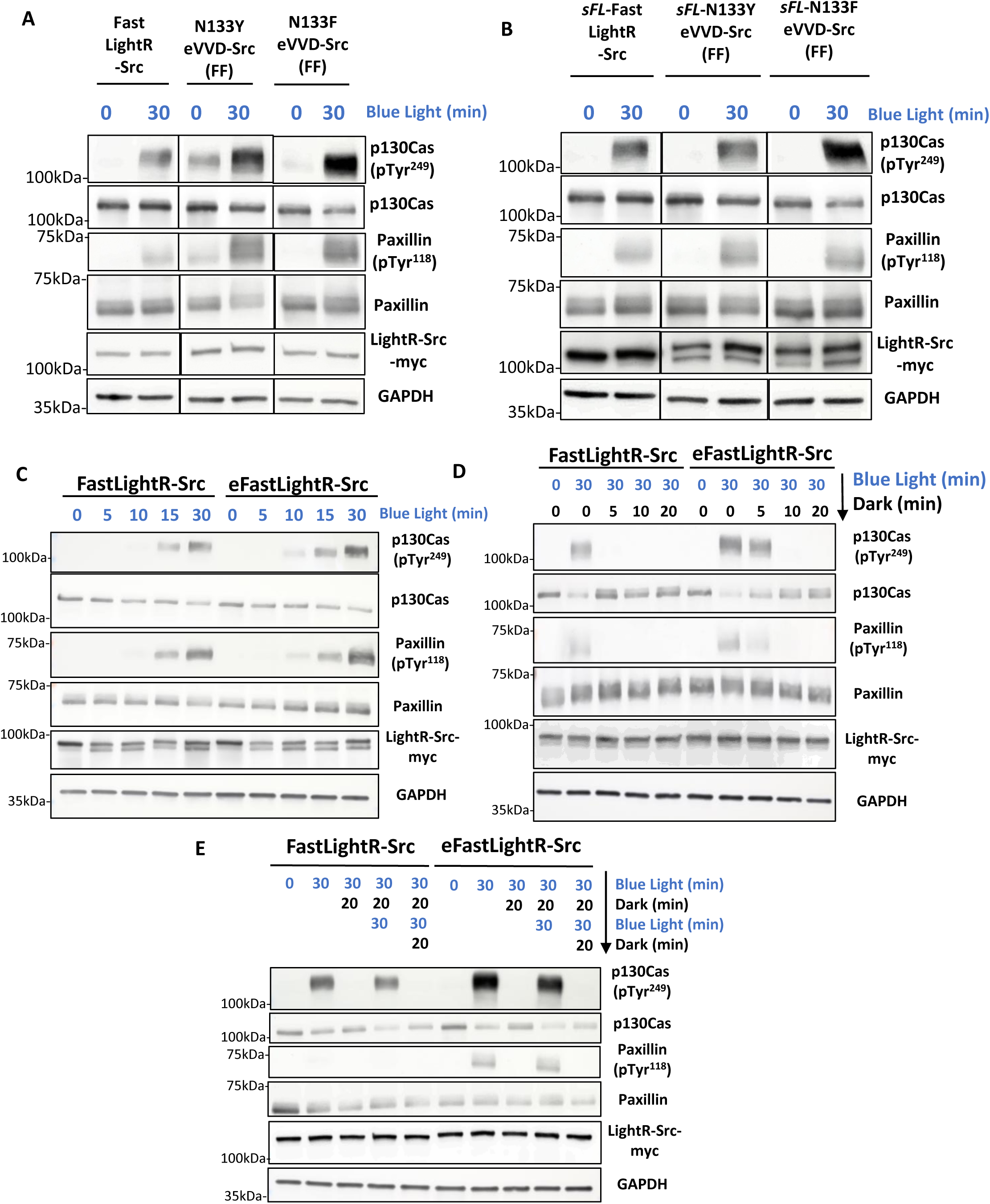
sFL linker in conjugation with stabilizing mutations improves dynamic range of FastLightR-Src regulation. Phosphorylation of endogenous p130 Cas and Paxillin, in response to activation of indicated Src constructs in living cells. (A-E) HEK293T cells transiently expressing the indicated Src constructs, bearing mCherry and myc tag at the C-terminus, were either kept in the dark or exposed to blue light for indicated periods of time. Cells, where dark time periods are indicated, were then placed in the dark for indicated time periods following illumination. Cell lysates were collected and probed for phosphorylation of Src substrates via Western Blot. (A) Comparison of activation between FastLightR-Src constructs bearing either original FastLightR domain or new versions comprised of indicated eVVD mutants. (B) Comparison of activation of different FastLightR-Src and/or eVVD mutant constructs bearing sFL inter-VVD linker. (C, D) Comparison of activation (C) and inactivation (D) kinetics for FastLightR-Src and eFastLightR-Src. (E) Repeated activation/inactivation cycles of FastLightR-Src and eFastLightR-Src. All experiments were repeated at least three times.

Our previous studies demonstrated that FastLightR-Src induces cell spreading upon illumination with blue light ^17^. We evaluated the ability of eFastLightR-Src to induce the same phenotype and compare the efficiency of activation with the previous FastLightR-Src construct. Our analysis revealed that upon illumination, eFastLightR-Src induces significantly faster and more robust spreading in HeLa cell (**Figure 6**). Overall, by combining stabilizing mutations with more rigid inter-VVD linker sFL we generated light-regulated allosteric switch that enables faster and stronger activation upon illumination while exhibiting fast inactivation when placed in the dark.

**Figure 6.**
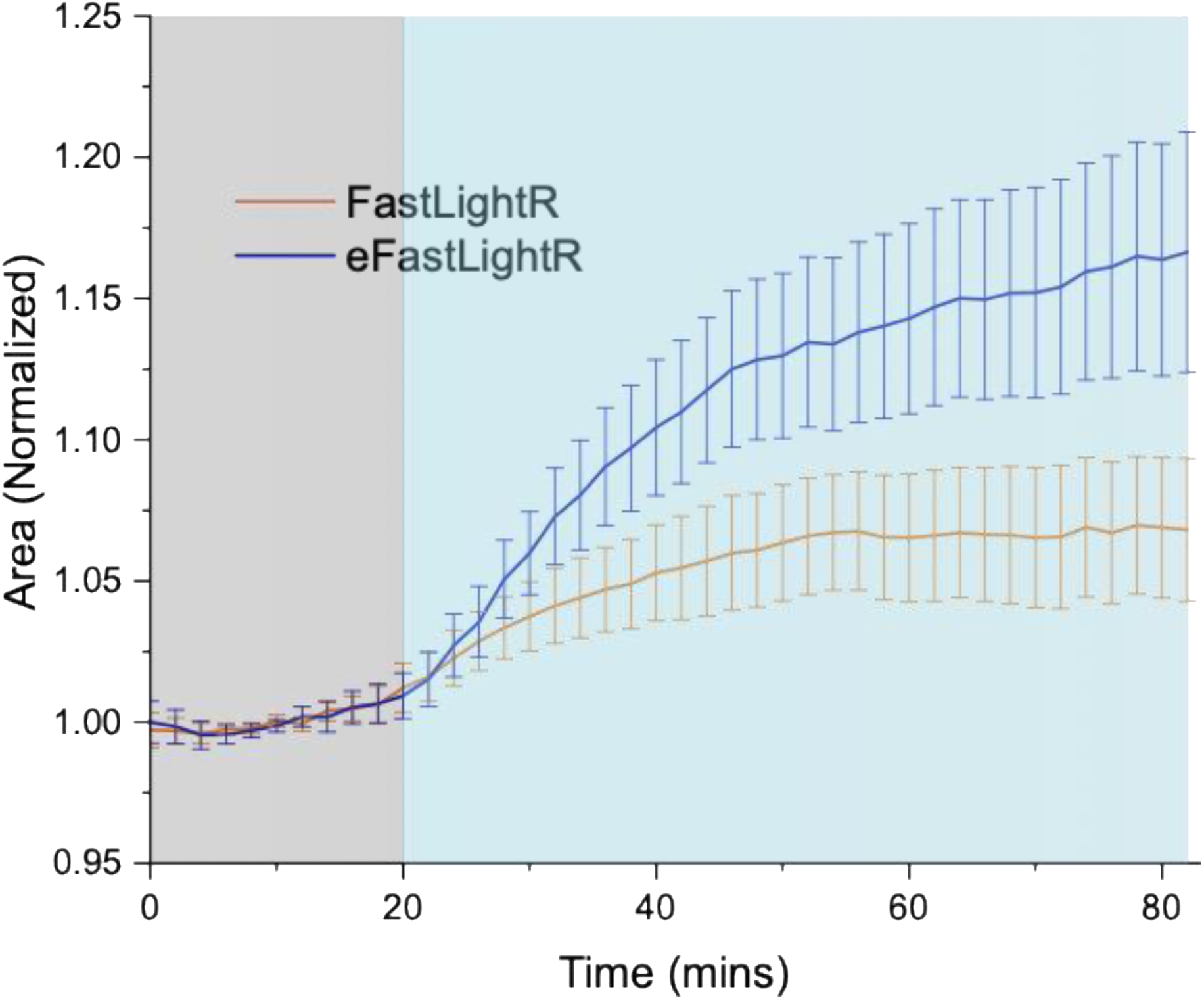
Regulation of cell morphology is enhanced with eFastLightR-Src: HeLa cells transiently co-expressing either FastLightR (N = 23 cells) or eFastLightR-Src (N = 19 cells) with stargazin-iRFP670 (plasma membrane marker) were imaged live every 2 minutes. Quantification of changes in cell area is shown before (grey box) and during (blue box) illumination with blue light. Graph represents mean ± 90% confidence intervals.

## Discussion

Our study aimed at the development of protein engineering approaches that will expand tunability and applicability of the LightR tool. We focused on modifications in two individual modules of LightR domain, photosensitive VVD domains and the linker connecting these domains. Mutations in the VVD domains that enhance protein stability in mammalian cells, strengthen their homodimerization, and reduce photoconversion rate from the lit state to the dark state consistently increased overall activity of LightR-Src. However, this was achieved at the expense of markedly increased activity of LightR-Src in the dark. Connecting VVD domains with a more rigid sFL linker significantly reduced leaky activity of LightR constructs generating a new module that can be utilized for tuning light-mediated regulation. Combined implementation of these two approaches incorporates their specific advantages and allows us to improve activation kinetics, achieve optimal dynamic range of activity, minimize leaky activity in the dark, and increase overall activity upon illumination.

Homodimerization of VVD domains occurs due to light-induced conformational changes in their structure. Spontaneous conversion of a lit state structure to a dark state can occur even during illumination and thus will temporarily break interaction between two VVD domains. In the context of LightR domain, this may lead to a reduced level of activity of the targeted protein. Conversely, spontaneous acquisition of the lit state in the dark may transiently cause VVD interaction favoring “closed” LightR domain conformation and thus result in dark leakiness of the targeted protein. Previously reported M135I/M165I double mutation dramatically slows down thermal reversion of lit conformation of VVD into dark state ^23^. Not surprisingly, introduction of these mutations into both VVD domains of LightR switch causes strong leakiness in the dark as even transient spontaneous interaction of the mutant VVD domains will become very stable (**Figure 1B**). Similarly, thermostable eVVD mutants may also possess the same property and thus result in the dark leakiness of LightR-mediated regulation (**Figure 5A**). Interestingly, N133F mutation in the eVVD caused reduced leakiness of LightR when compared with to N133Y mutation (**Figure 5A**). This is in agreement with the previously reported faster reversal of N133F to the dark state ^1^. However, how these mutations mediate enhancement or reduction of cysteine adduct state in VVD remains unclear. Crystal structure of lit VVD dimer suggests that Tyr or Phe at position 133 should interact with adenine portion of the flavin adenine dinucleotide, potentially making a pi-pi stacking interaction. How this interaction enhances the photoadduct formation and why Phe would promote faster reversal to the dark state remains to be elucidated.

Spider silk and ferredoxin-like proteins have been used as linkers in several applications where their spring-like behavior was utilized to measure tension applied to a specific protein of interest in living cells ^18,19^. This unique property spring-like linkers also suggests that in the collapsed form their N- and C-termini will stay in a more rigid position when compared to highly flexible Gly-Pro-Gly repeat linker. However, their ability to “stretch” also indicates that they could provide sufficient flexibility that allows to bring N- and C-termini close. Our modeling supports this hypothesis (**Figure 2**) and our biochemical analysis shows that sFL linker has the optimal properties that diminish leaky activity of LightR domain in the dark and allows for robust activation upon illumination (**Figure 3A**). The inability of SSL linker to reduce leaky activity can be explained by the predicted tendency to favor shorter distances between N- and C-termini (**Figure 2C**). Worm-like chain proteins are known for fluctuating between fully extended, more rigid, and collapsed structures ^20^. However, previous studies analyzed significantly longer peptides than the linker used in our study ^17^. Our analysis suggested that shortened worm-like chain linker could still acquire an extended and a collapsed form (**Figure 2C**). However, when inserted into LightR, it failed to reduce dark leakiness sufficiently (**Figure 3A**). This is potentially due to inability of the shortened worm-like chain linker to hold rigid structure when inserted in the LightR domain. A longer worm-like chain linker may provide more rigidity.

## Materials and Methods

### Antibodies, Reagents, and Cell Lines

The following antibodies were used: anti-Myc (Millipore, cat. no. 05–724), anti-paxillin (Fisher Scientific, cat. no. BDB612405), anti-phospho-paxillin (Tyr^118^) (Invitrogen, cat. no. 44–722G), anti-p130cas (BD biosciences, cat. no. 610271), anti-phospho-p130cas (Tyr^249^) (BD biosciences, cat. no. 558401) The following cell lines were used: HeLa cells (ATCC, cat. No. CCL- 2), Human embryonic kidney HEK293T cells (ATCC, cat. no. CRL-3216). All cell lines were cultured at 37°C and 5% carbon dioxide in DMEM medium supplemented with 10% v/v Fetal Bovine Serum, 1% v/v GlutaMAX^TM^, and 1% non-essential amino acids (cat. no. 11140-050). All experiments were performed with cells grown for less than 20 passages after thawing. All cells were tested negative for *Mycoplasma* contamination. Cell line identity was confirmed by the supplier using STR analysis.

The following reagents were used: 4X Laemmli Sample buffer (BIO-RAD, cat. no. 161–0747), 2-Mercaptoethanol (Fisher Chemical, cat. no. 60-24-2), trypsin (Promega, cat. no. V5113), accutase (Corning, cat. no. 10210-214), Fetal Bovine Serum (Omega Scientific, cat. no. FB-01), and , GlutaMAX^TM^ (ThermoFisher, cat. no. 35050061), non-essential amino acids (Gibco^TM^, cat. no. 11140-050), JetOptimus transfection reagent (Polyplus, cat. no. 101000006). PEI HTS transfection reagent (Polyplus, cat. no. 101000053). Coverslips for live imaging were purchased from ThermoFisher (cat. no. 25CIR-1.5).

### Molecular biology and expression plasmids

LightR-Src-mCherry-myc, FastLightR-Src-mCherry-myc (F_1_/F_1_), and LightR-Src-mCherry-myc (H_1_/H_1_) were generated by Mark Shaaya as described in Shaaya et. al ^17^. Using site direct mutagenesis the High1 mutations (M135I, M165I) were introduced in the C-terminal VVD in the LightR switch domain. Replacement of the inter-VVD linkers were performed through generation of a mega primer containing the linker sequence followed by site directed mutagenesis, as previously described ^24^. The original worm-like chain helix is 10-30 nm in length, which could prevent a closed conformation of light activated LightR switch. To avoid this, we truncated WLC linker (22 amino acids). The truncated sequence retains the charged Glu repeats followed by Arg or Lys repeats with stochastic break points to enable transitions between the extended and collapsed conformations. To produce the thermostabilized enhanced VVDs (eVVD) we introduced the following mutations via site directed mutagenesis: T69L, Y94E, S99A, N100R, A101H, N133Y/F, R136K, and M179I. These mutations and linker changes were performed in LightR-Src-mCherry-myc.

### Amino Acid sequences of inter-VVD linkers

Original GSG Linker: GGSGGSGGSGGSGGGSGGSGGS ;

Spider-Silk Linker: GGPGGAGPGGAGPGGAGPGGAG ; Worm-Like Chain Linker: GEEEEKKKQQEEEAERLRRIQG ;

Ferredoxin-Like Linker: GEFDIRFRTDDDEQFEKVLKEMHRRARKDAG

### Transient expression of constructs in HEK293T cells and activity assessment

For expression of the LightR constructs, HEK293T cells were grown to 70% confluency on a plastic dish coated with 1:200 poly-L-lysine and were transfected with JetOptimus transfection reagent following manufacturer’s protocol. Cells were continuously stimulated with a blue light using the HQRP LED Plant Grow Panel Lamp System (3 mW/cm^2^, 465 nm wavelength) for the indicated periods of time. Control cells were kept in the dark. Cells were then lysed using the lysis buffer (20 mM HEPES-KOH, pH 7.8, 50 mM KCl, 1 mM EGTA, 1% NP40, 1 mM NaF, 0.2 mM Na3VO4, aprotinin 16 µg/ml, and Leupeptin hemisulfate 3.2 µg/mL). Lysates were centrifuged at 4000 rpm, 4°C, for 10 min, and the cleared lysates were incubated with 4X Laemmli sample buffer with 10% 2-Mercaptoethanol at 95-100°C for 5minutes, proteins were resolved via sodium dodecyl sulfate-polyacrylimide gel electrophoresis (SDS/PAGE) and immunoblotted. Blots were developed using enhanced chemiluminescence.

### In silico analysis of inter-VVD linkers

Theoretical structures of the inter-VVD linkers were produced using Chimera structure builder ^25^. These were then minimized using steepest descent, equilibrated at 310 K and simulated for 1000 frames in Chimera. Three starting structures for each linker were then selected from frames within these simulations. All structures were energy minimized in NAMD (version 2.13) using a steepest descent gradient for 1000 steps ^27^ These structures were then equilibrated for 0.5 ns at increasing temperature from 60 K to 310 K, then for an additional 1 ns at 310 K, with generalized Born implicit solvent with an ion concentration of 0.15 M. MD simulations were carried out, using these same conditions, over 45 ns for each of the three starting structures across all linkers. RMSF was averaged and visualized from each of these simulations using VMD (version 1.9.3).

### GST Paxillin Purification

N-terminal fragment of paxillin was purified following previously described procedure ^28^. Briefly, GST-paxillinN-C3 construct was expressed in DH5α bacteria cells following induction with 0.1 mM Isopropyl β-D-1-thiogalactopyranoside for 4 hr. Bacterial pellet was resuspended in 30 ml of TETN buffer (20 mM TRIS pH 8, 100 mM NaCl, 1 mM EDTA, 0.5% Triton X100) and lysed by sonication. GST-paxillinN-C3 was purified from the cleared lysates by affinity chromatography using Glutathione Sepharose following previously described protocol ^28^

### *In vitro* Kinase Assay

A modification of a previously described protocol for in vitro kinase assay was used in all assays^24^. Briefly, Src kinase constructs bearing an mCherry and a myc tandem tags at the C-terminus were transiently overexpressed in 293T cells. Cells were exposed to continuous blue light (3 mW/cm^2^, 465 nm wavelength) for the indicated times or kept in the dark. Lysates were then collected under red light illumination (to prevent activation of LightR) using the lysis buffer (20 mM HEPES-KOH, pH 7.8, 50 mM KCl, 1 mM EGTA, 1% NP40, 1 mM NaF, 0.2 mM Na3VO4, aprotinin 16 µg/ml, and Leupeptin hemisulfate 3.2 µg/mL). Lysates were centrifuged at 4000 rpm, 4°C, for 10 min, and the cleared lysates were incubated with ProteinG sepharose beads conjugated with the anti-myc antibody (4A6 from Millipore-Sigma) for 1.5 hr at 4°C. Beads were then washed with wash buffer (20 mM Hepes-KOH, pH 7.8, 100 mM NaCl, 50 mM KCl, 1 nM EGTA, 1% NP40) and then with kinase reaction buffer (25 mM HEPES, pH 7.5, 5 M MgCl_2_, 0.5 mM EGTA, 0.005% BRIJ-35). The beads were resuspended in kinase reaction buffer and incubated with 0.1 mM ATP and indicated concentrations of purified N-terminal fragment of paxillin (GST-paxillinN-C3) at 37°C for indicated timepoints. The reaction was terminated by adding 4X Laemmli sample buffer with 10% v/v 2-Mercaptoethanol then incubating at 95-100°C for 5 min. The phosphorylation of paxillin was examined by western blotting. Initial reaction velocities were determined by quantifying phosphorylated paxillin using western blot densitometry analysis. The velocity values were plotted as a function of substrate concentration, and kinetic parameters (Km and Vmax) were obtained by nonlinear regression fitting to the Michaelis–Menten equation using GraphPad Prism.

### Live cell imaging and analysis

HeLa cells were grown to 60% confluency in a 60 mm dish and transfected with 4µL of PEI HTS and 2 µg of total DNA and incubated overnight. Cells were co-transfected with LightR-Src andstargazin-iRFP670 as a membrane marker. Two days post transfection, cells were seeded at 30-40% confluency on fibronectin-coated (10 mg/L) glass coverslips for 2-3 hours prior to imaging and imaged in Leibovitz L-15 imaging media. Cells were imaged live at 37 °C in an open heated chamber (Warner Instruments) using Olympus UPlanSAPO 40x (oil, N.A. 1.25) objective on Olympus IX-83 microscope controlled by Metamorph software. Time lapse images were acquired every 2 minutes over the duration of the imaging experiment. Light activation was performed using a blue LED ring light (BoliOptics cat. no. ML46241324) attached to the microscope condenser and controlled by MetaMorph software. Cells were illuminated for 40 seconds per minute during the illumination phase of each experiment.

Epifluorescence images of stargazin-iRFP670 were used to analyze changes in cell morphology. The binary mask of the images was created using MovThresh2014 ^29^ software package in Matlab (version 2017a). Cell area, in pixels, was calculated in ImageJ for each timepoint. To determine the area change for each cell, the area at each time point was normalized to the average area of the cell during the time prior to stimulation with blue light. The average of these normalized values and 90% confidence interval were then calculated.

## Notes

### Competing Interest Statement

The authors have declared no competing interest.

## References

1. Benedetti, L., Marvin, J. S., Falahati, H., Guillén-Samander, A., Looger, L. L., and De Camilli, P. (2020) Optimized vivid-derived magnets photodimerizers for subcellular optogenetics in mammalian cells. Elife. 9, 1–49

2. Goglia, A. G., Wilson, M. Z., Jena, S. G., Silbert, J., Basta, L. P., Devenport, D., and Toettcher, J. E. (2020) A Live-Cell Screen for Altered Erk Dynamics Reveals Principles of Proliferative Control. Cell Syst. 10, 240–253.e6

3. Guntas, G., Hallett, R. A., Zimmerman, S. P., Williams, T., Yumerefendi, H., Bear, J. E., and Kuhlman, B. (2015) Engineering an improved light-induced dimer (iLID) for controlling the localization and activity of signaling proteins. Proc. Natl. Acad. Sci. U. S. A. 112, 112–117

4. Kawano, F., Suzuki, H., Furuya, A., and Sato, M. (2015) Engineered pairs of distinct photoswitches for optogenetic control of cellular proteins. Nature Communications *2015 6:1.* 6, 1–8

5. Mühlhäuser, W. W. D., Weber, W., and Radziwill, G. (2019) OpEn-Tag-A Customizable Optogenetic Toolbox To Dissect Subcellular Signaling. ACS Synth. Biol. 8, 1679–1684

6. Idevall-Hagren, O., Dickson, E. J., Hille, B., Toomre, D. K., and De Camilli, P. (2012) Optogenetic control of phosphoinositide metabolism. Proc. Natl. Acad. Sci. U. S. A. 109, E2316–E2323

7. Shaaya, M., Fauser, J., and Karginov, A. V. (2020) Optogenetics: The Art of Illuminating Complex Signaling Pathways. Physiology. 36, 52

8. Dagliyan, O., Krokhotin, A., Ozkan-Dagliyan, I., Deiters, A., Der, C. J., Hahn, K. M., and Dokholyan, N. V. (2018) Computational design of chemogenetic and optogenetic split proteins. Nature Communications *2018 9:1.* 9, 1–8

9. Bunnag, N., Tan, Q. H., Kaur, P., Ramamoorthy, A., Sung, I. C. H., Lusk, J., and Tolwinski, N. S. (2020) An Optogenetic Method to Study Signal Transduction in Intestinal Stem Cell Homeostasis. J. Mol. Biol. 432, 3159–3176

10. Nguyen, M. K., Kim, C. Y., Kim, J. M., Park, B. O., Lee, S., Park, H., and Heo, W. Do (2016) Optogenetic oligomerization of Rab GTPases regulates intracellular membrane trafficking. Nat. Chem. Biol. 12, 431–436

11. Dagliyan, O., Dokholyan, N. V., and Hahn, K. M. (2019) Engineering proteins for allosteric control by light or ligands. Nat. Protoc. 14, 1863

12. Hongdusit, A., and Fox, J. M. (2021) Optogenetic Analysis of Allosteric Control in Protein Tyrosine Phosphatases. Biochemistry. 60, 254–258

13. Winkler, A., Barends, T. R. M., Udvarhelyi, A., Lenherr-Frey, D., Lomb, L., Menzel, A., and Schlichting, I. (2015) Structural details of light activation of the LOV2-based photoswitch PA-Rac1. ACS Chem. Biol. 10, 502–509

14. Strickland, D., Moffat, K., and Sosnick, T. R. (2008) Light-activated DNA binding in a designed allosteric protein. Proc. Natl. Acad. Sci. U. S. A. 105, 10709–10714

15. Hongdusit, A., Liechty, E. T., and Fox, J. M. (2022) Analysis of Three Architectures for Controlling PTP1B with Light. ACS Synth. Biol. 11, 61–68

16. Hongdusit, A., Zwart, P. H., Sankaran, B., and Fox, J. M. (2020) Minimally disruptive optical control of protein tyrosine phosphatase 1B. Nature Communications *2020 11:1.* 11, 1–11

17. Shaaya, M., Fauser, J., Zhurikhina, A., Conage-Pough, J. E., Huyot, V., Brennan, M., Flower, C. T., Matsche, J., Khan, S., Natarajan, V., Rehman, J., Kota, P., White, F. M., Tsygankov, D., and Karginov, A. V. (2020) Light-regulated allosteric switch enables temporal and subcellular control of enzyme activity. Elife. 9, 1–73

18. Grashoff, C., Hoffman, B. D., Brenner, M. D., Zhou, R., Parsons, M., Yang, M. T., McLean, M. A., Sligar, S. G., Chen, C. S., Ha, T., and Schwartz, M. A. (2010) Measuring mechanical tension across vinculin reveals regulation of focal adhesion dynamics. Nature *2010 466*:7303. 466, 263–266

19. Ringer, P., Weißl, A., Cost, A. L., Freikamp, A., Sabass, B., Mehlich, A., Tramier, M., Rief, M., and Grashoff, C. (2017) Multiplexing molecular tension sensors reveals piconewton force gradient across talin-1. Nat. Methods. 14, 1090–1096

20. Sivaramakrishnan, S., and Spudich, J. A. (2011) Systematic control of protein interaction using a modular ER/K α-helix linker. Proc. Natl. Acad. Sci. U. S. A. 108, 20467–20472

21. Zoltowski, B. D., Vaccaro, B., and Crane, B. R. (2009) Mechanism-based tuning of a LOV domain photoreceptor. Nat. Chem. Biol. 5, 827–834

22. Karginov, A. V., and Hahn, K. M. (2011) Allosteric activation of kinases: Design and application of RapR kinases. Curr. Protoc. Cell Biol. **CHAPTER**, Unit

23. Pettersen, E. F., Goddard, T. D., Huang, C. C., Couch, G. S., Greenblatt, D. M., Meng, E. C., and Ferrin, T. E. (2004) UCSF Chimera--a visualization system for exploratory research and analysis. J. Comput. Chem. 25, 1605–1612

24. Šali, A., and Blundell, T. L. (1993) Comparative protein modelling by satisfaction of spatial restraints. J. Mol. Biol. 234, 779–815

25. Phillips, J. C., Braun, R., Wang, W., Gumbart, J., Tajkhorshid, E., Villa, E., Chipot, C., Skeel, R. D., Kalé, L., and Schulten, K. (2005) Scalable molecular dynamics with NAMD. J. Comput. Chem. 26, 1781–1802

26. Lyons, P. D., Dunty, J. M., Schaefer, E. M., and Schaller, M. D. (2001) Inhibition of the Catalytic Activity of Cell Adhesion Kinase β by Protein-tyrosine Phosphatase-PEST-mediated Dephosphorylation. Journal of Biological Chemistry. 276, 24422–24431

27. Tsygankov, D., Bilancia, C. G., Vitriol, E. A., Hahn, K. M., Peifer, M., and Elston, T. C. (2014) CellGeo: A computational platform for the analysis of shape changes in cells with complex geometries. Journal of Cell Biology. 204, 443–460

