## supplementary figures and figure legend for "Tuning a light-regulated allosteric switch for enhanced temporal control of protein activity"

### Supplementary Figure 1

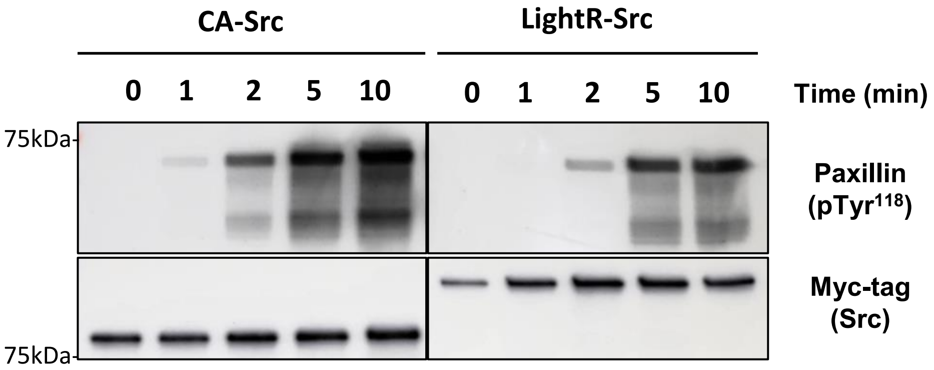

### Supplementary Figure 2

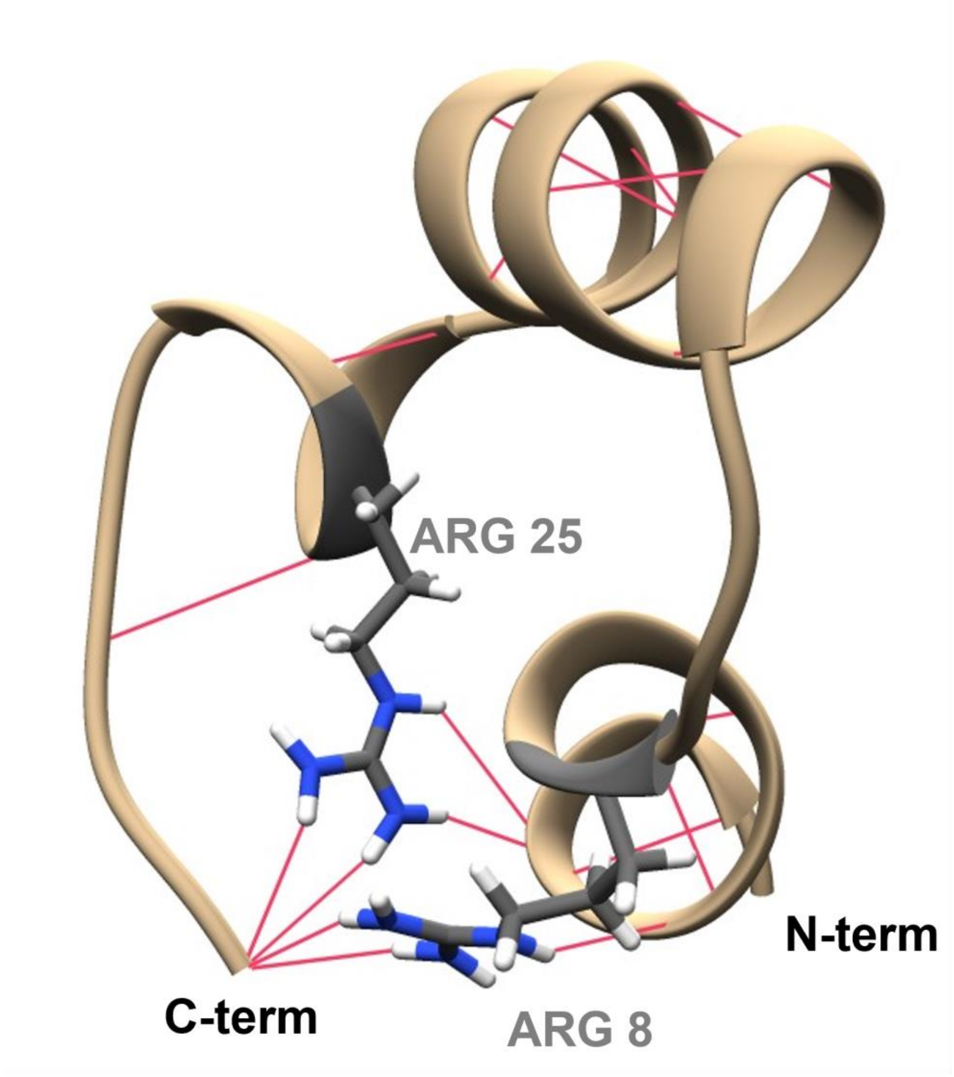

### Supplementary Figure 3

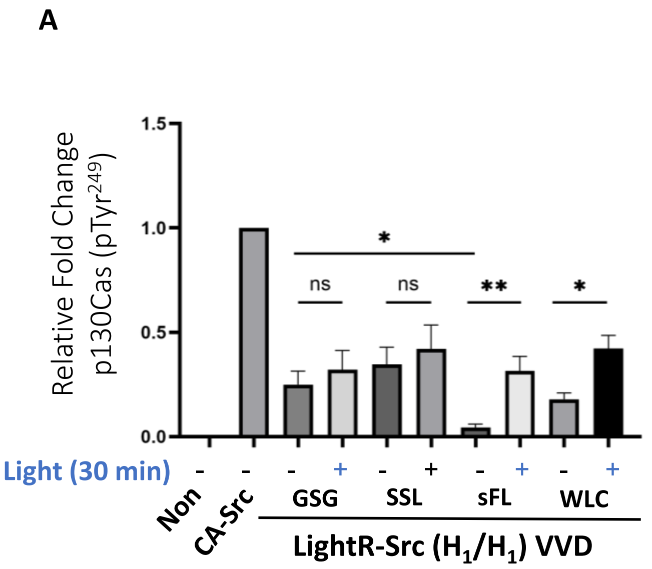


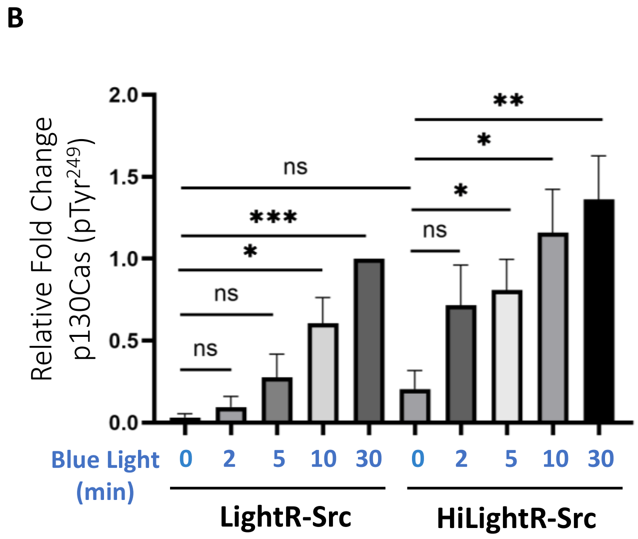


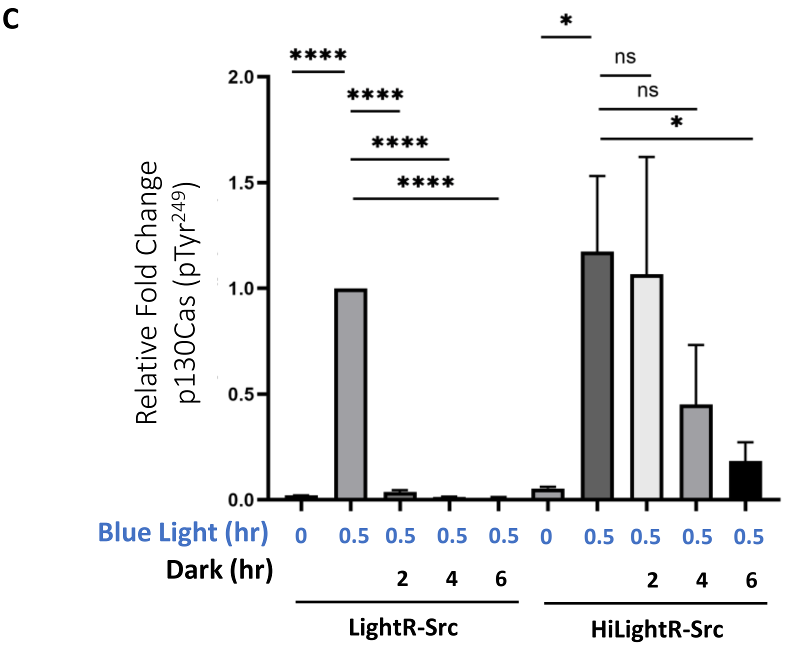

### Supplementary Figure 4

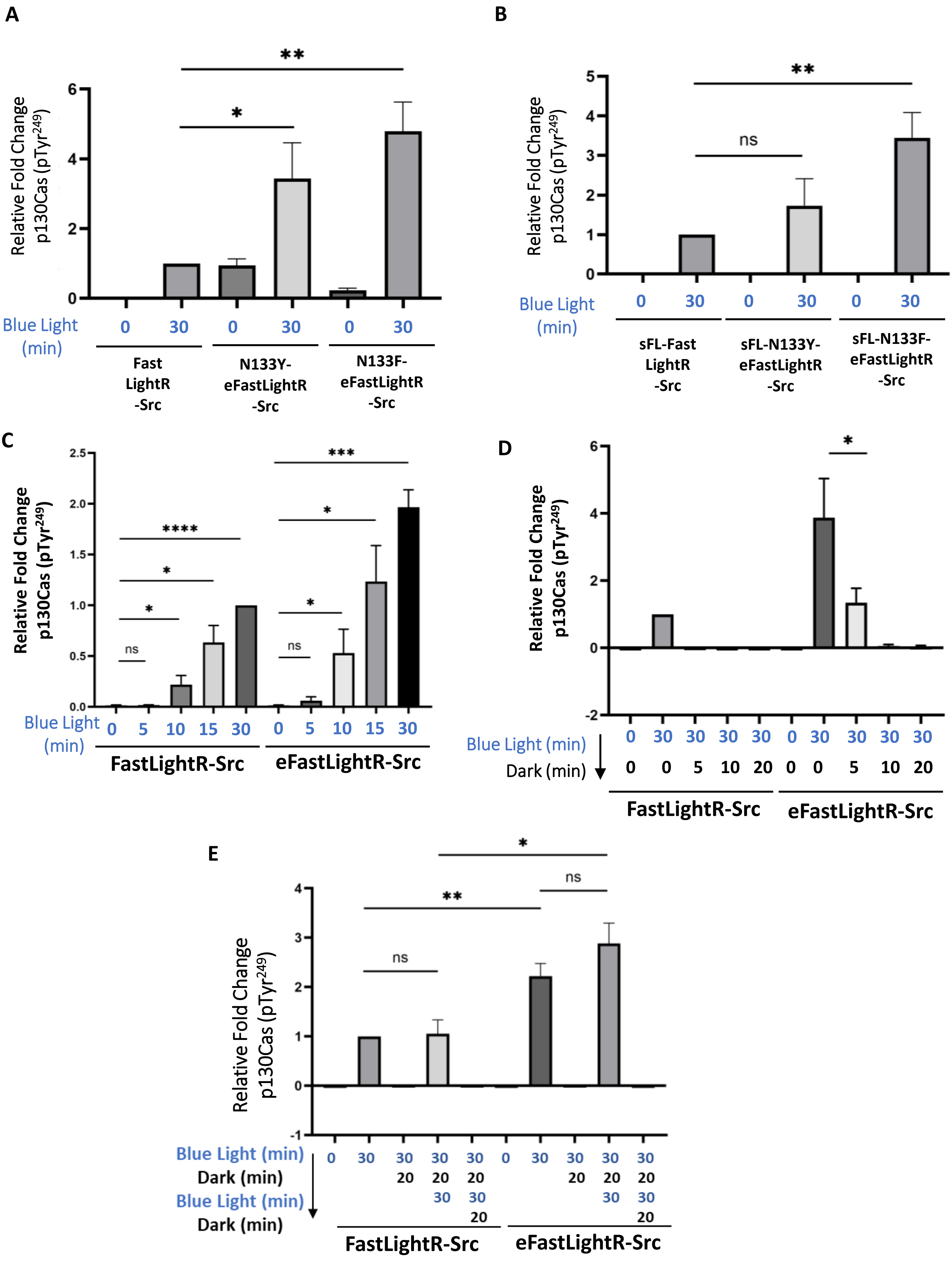

**Supplementary Figure Legends:**

**Supplementary Figure 1. Analysis of LightR-Src and constitutively active (CA) Src activity using *in vitro* assay.** 293T cells transiently expressing the indicated Src constructs bearing mCherry and myc tags at the C-terminus were exposed to continuous blue light for 30 minutes. Src constructs were then immunoprecipitated and their ability to phosphorylate purified N-terminal fragment of paxillin was assessed by in vitro kinase assay.

**Supplementary Figure 2: Ferredoxin-like linker hydrogen bonding network.** Representative structure of ferredoxin-like linker from molecular dynamics simulations. Three starting structures were generated, minimized, and equilibrated in Chimera before performing MD simulations at 310 K in an implicit solvent field using NAMD for 45 ns. Hydrogen bonds indicated in pink, were mapped in averaged structures from the MD simulation using Chimera. Arg8 and Arg25 are highlighted in grey to show important interactions.

**Supplementary Figure 3: sFL inter-VVD linker in combination with M135I/M165I mutations improve activation of LightR-Src.** (A-C) Quantification of p130Cas phosphorylation described in **Figure 3A** (A), **3B** (B), and **3E** (C). Phosphorylation levels were measured as relative ratio of phospho(Tyr 249) p130Cas level to total p130Cas levels in each group normalized to the value for CA-Src (A) or LightR-Src sample at 30 minutes of activation (B, C). All experiments were repeated at least three times with similar results. All analyses graphs are shown as mean ± standard errors of mean (SEM) and statistical differences were examined by unpaired Student's t-test. NS: not significant ($p>0.05$) ; * : significant ($p<0.05$) ; *** : significant ($p<0.001$) ; **** : significant ($p<0.0001$).

**Supplementary Figure 4. sFL linker in conjugation with stabilizing mutations improves dynamic range of FastLightR-Src regulation.** Quantification of p130Cas phosphorylation from experiments described in Figure 4A (A), 4B (B), 4C (C), 4D (D) and 4E (E). Phosphorylation levels were measured as relative ratio of phospho(Tyr 249) p130Cas level to total p130Cas levels in each group normalized to the value for FastLightR-Src sample at 30 minutes of activation. All experiments were repeated at least three times. All analyses graphs are shown as mean ± standard errors of mean (SEM) and statistical differences were examined by unpaired Student's t-test. NS: not significant ($p>0.05$) ; * : significant ($p<0.05$) ; *** : significant ($p<0.001$) ; **** : significant ($p<0.0001$).
